# SUMIC: A Simple Ultrafast Multicolor Immunolabelling and Clearing Approach for Whole-Organ and Large Tissue 3D Imaging

**DOI:** 10.1101/2021.01.20.427385

**Authors:** Lincoln Biswas, Junyu Chen, Jessica De Angelis, Alexandros Chatzis, Jagdeep Nanchahal, Michael L. Dustin, Saravana K. Ramasamy, Anjali P. Kusumbe

**Affiliations:** Tissue and Tumor Microenvironments Group, University of Oxford, Kennedy Institute of Rheumatology, NDORMS, Oxford, OX3 7FY, UK; University of Oxford, Kennedy Institute of Rheumatology, Oxford, OX3 7FY, UK; Institute of Clinical Sciences, Imperial College London, London W12 0NN, UK; MRC London Institute of Medical Sciences, Imperial College London, London W12 0NN, UK; Department of Prosthodontics, State Key Laboratory of Oral Diseases, West China Hospital of Stomatology, Sichuan University, Chengdu 610041, China

**Keywords:** 3D imaging, blood vessels, lymphatic vessels, tissue clearing, light sheet

## Abstract

High-resolution whole-organ imaging of cleared tissues captures cellular and molecular insights within the intact tissue and tumour microenvironments. However, current immunolabelling and clearing methods are complicated and time-consuming; extending to several weeks. Here, we developed **<u>S</u>**imple **<u>U</u>**ltrafast **<u>M</u>**ulticolor **<u>I</u>**mmunolabelling and **<u>C</u>**learing or **SUMIC**, a method that enables multicolor immunolabelling and clearing of whole murine organs and human tissues within 2 to 2.5 days. Moreover, SUMIC is simple, robust, non-hazardous and versatile comprising antigen retrieval, permeabilization, collagenase-based digestion, immunolabelling, dehydration, and clearing. SUMIC permits quantitative and singlecell resolution analysis and detection of rare cells in whole organs, for example, round αSMA positive cells in the thymus. Upon volumetric imaging, SUMIC-processed samples retain normal tissue architecture and can be used for paraffin-embedding and histology. We employed the SUMIC method for whole-organ mapping of lymphatic vessels across different ages and organs. This analysis revealed the expansion of lymphatic vessels in endocrine tissues but not in any other organs with aging. Hence, SUMIC will accelerate discoveries compared to other whole organ imaging pipelines.

## Introduction

Advanced 3D imaging approaches provide critical insights into the fundamentals of highly complex biological processes during organ development, regeneration, repair and pathology (*1–7*). To advance our understanding of such processes, it is vital to generate accurate cellular, and molecular cartography of healthy, aged, diseased and malignant tissues is vital (*8–11*). The 3D analysis of intact biological specimens reveals essential structural and spatial information necessary for the understanding of the crosstalk among various cell types in the tissue context (*9, 12–16*). Moreover, deciphering the organization of the nervous and vascular systems relies on a whole organ approach (*12, 17–20*). This treasure trove of information comes at the expense of detailed and complicated requirements for time-consuming sample processing, imaging, data storage, analysis and interpretation, which can be met by a range of recent technical developments.

The recent development of tissue clearing, light sheet microscopy and large dataset analysis has made it possible to image large volumes at single-cell resolution (*13, 21–23*). However, current protocols for immunostaining and whole organ clearing are often time-consuming, elaborate, expensive, and organ-specific (*24–32*). The time-consuming nature of the whole organ and large tissue imaging is the main obstacle to overcome before efficiently applying this crucial technique to understand biological complexity and interrogating cellular interactions within tissues and organs. Other disadvantages include the requirement for transgenic mice that express fluorescent reporters and the use of toxic or corrosive chemicals that might be detrimental either to the microscope or researcher’s well-being (*11, 33–35*).

Therefore, it is essential to develop a rapid method for whole organ and large tissue multicolor immunostaining coupled with a technique applicable for diverse tissues while preserving endogenous fluorescence. Here we devised a new robust and straightforward method-**<u>S</u>**imple **<u>U</u>**ltrafast **<u>M</u>**ulticolor **<u>I</u>**mmunolabelling and **<u>C</u>**learing or SUMIC which permits immunostaining and clearing of multiple whole organs and large tissues in a record time of 2.5 days, and also overcomes other limitations of current methods.

## Results

### SUMIC: an ultrafast immunostaining and clearing method for murine organs and human tissues

In SUMIC we set out to develop a platform that is able to attain a multicolor immunolabelling together with excellent cellular morphology, tissue architecture and transparency across multiple murine organs and human tissues. Each of the 10 steps were optimized to enable completion in the shortest possible time (Fig. 1). First, we combined paraformaldehyde and glutaraldehyde for rapid fixation (Step 1). Samples were bleached with hydrogen peroxide in methanol to eliminate heme and other persistent chromophores (Step 2). Urea facilitated antigen retrieval, and Triton X-100 increased the penetration depth of antibodies into tissues while maintaining tissue architecture (Step 3). Collagenase-digestion of the extracellular matrix significantly improved the penetration rate of the antibodies while keeping the structure of the tissues intact (Step 4). To obtain good staining, only antibodies labelled with small, bright fluorochromes with high stability and good tissue penetrance, such as Alexa Fluor dyes, were used (Step 5). Triton X 100, a mild detergent was included for blocking, staining and washing the organs during this step. Incubations with antibodies and their washings were performed at 37°C to reduce the time requirement further. Dehydration is a crucial step that precedes tissue clearing and RI matching (Step 6). Dehydration using isopropanol preserved all fluorochromes and resulted in less tissue shrinkage compared to ethanol. Ethyl cinnamate (ethyl 3-phenyl-2-propenoate; ECi) is non-toxic and has been utilized for tissue clearing (*11*). Poly(ethylene glycol) methyl ether methacrylate (PEGM) has also been used as a component of tissue clearing solutions (*26*). For SUMIC ECi (80%) and PEGM (20%) were combined. This combination of ECi and PEGM cleared all the tested organs and tissues within 2 hours (Step 7). ECi was used for RI matching for imaging (Step 8). Starting from collecting the organs to image acquisition, SUMIC protocol for multicolor antibody labelling and tissue clearing required up to 2.5 days with conjugated antibodies (Fig. 1). SUMIC also includes a rendering and analysis pipeline (Step 9).

**Fig. 1.**
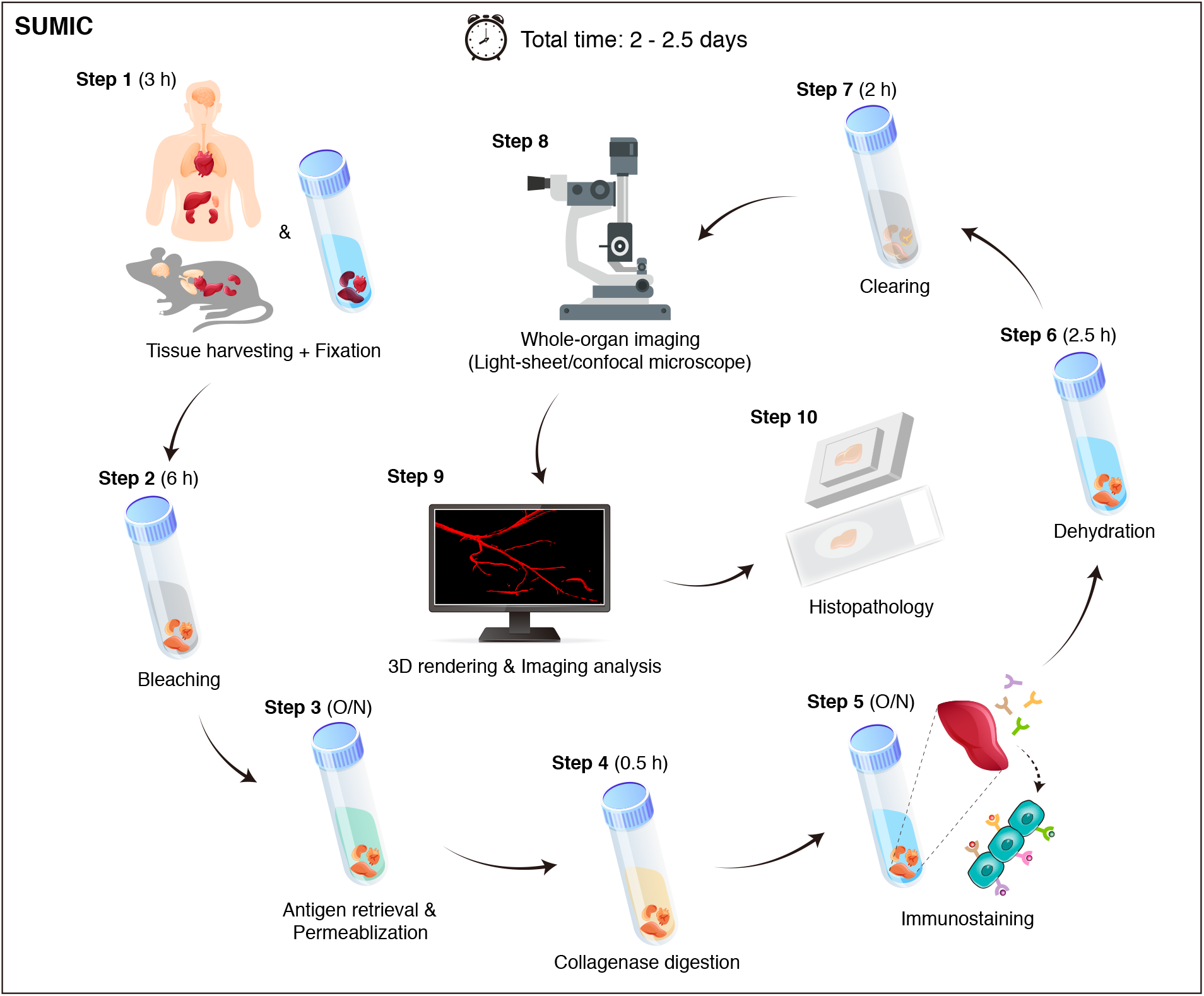
Whole-organ 3D imaging using the simple ultrafast multicolor immunolabelling and clearing (SUMIC) method. Schematic illustration of a simple ultrafast multicolor immunolabelling and clearing (SUMIC) method for whole-organ imaging. After tissue harvesting and fixation (step 1), samples then go through bleaching (step 2, optional), antigen retrieval and permeabilization (step 3), collagenase digestion (step 4), immunostaining (step 5), dehydration (step 6) and clearing (step 7). Whole-organ images of cleared organs are acquired using a light-sheet or confocal microscope (step 8), and Z-stacks are subsequently 3D rendered and analyzed (step 9). After imaging, samples can also be used for histopathology (step 10, optional). The time shown for each step is the maximum time required to perform that step. The total time of the SUMIC is around 2.5 days. O/N, overnight.

Finally, after SUMIC, the clarified tissues could be sectioned histopathology (Step 10).

We have successfully validated SUMIC across multiple murine organs, including lung, heart, kidney, liver, gut, spleen, thymus, prostate gland, seminal vesicle and endocrine glands (Fig. 2, Fig. 3, Fig. S1 and Movie 1–6). Notably, a few organs, such as liver, demonstrated a slightly brownish yellow colour even after the completion of the clearing process (Fig. 2D); however, this had minimal impact on the penetration depth of the light sheet. We selected ~40 of antibodies to test and validate SUMIC method across multiple organs and tissues. We selected antibodies for the cell and matrix molecules expressed across different organs. The cell markers defining vascular and tissue microenvironments, including endothelial cells, pericytes, mesenchymal cells and immune cells, were used. Endothelial cell markers; Endomucin (Emcn), Endoglin (CD105), CD31 also known as PECAM1 and CD102 (ICAM2); a-SMA staining was used for artery quantifications, and pericytes were identified by PDGFRβ and NG2. Lymphatic vessels were analyzed by LYVE1 and Prox1. We provide full information for each of these antibodies and their working conditions (Table S1). With these cell surface markers, SUMIC enabled 3D imaging of blood vessels and associated niches across several organs and endocrine glands (Fig. 2, Fig. 3, Fig. S1 and Movie 1–6). Thus, the SUMIC method provided the flexibility of usage across multiple organs and tissues. Troubleshooting tips relevant to the SUMIC method can be found in Table S2, and the methods section of the paper. In addition to the murine organs, SUMIC permitted ultrafast immunostaining and clearing of human tissues (Fig. 4, A and B).

**Fig. 2.**
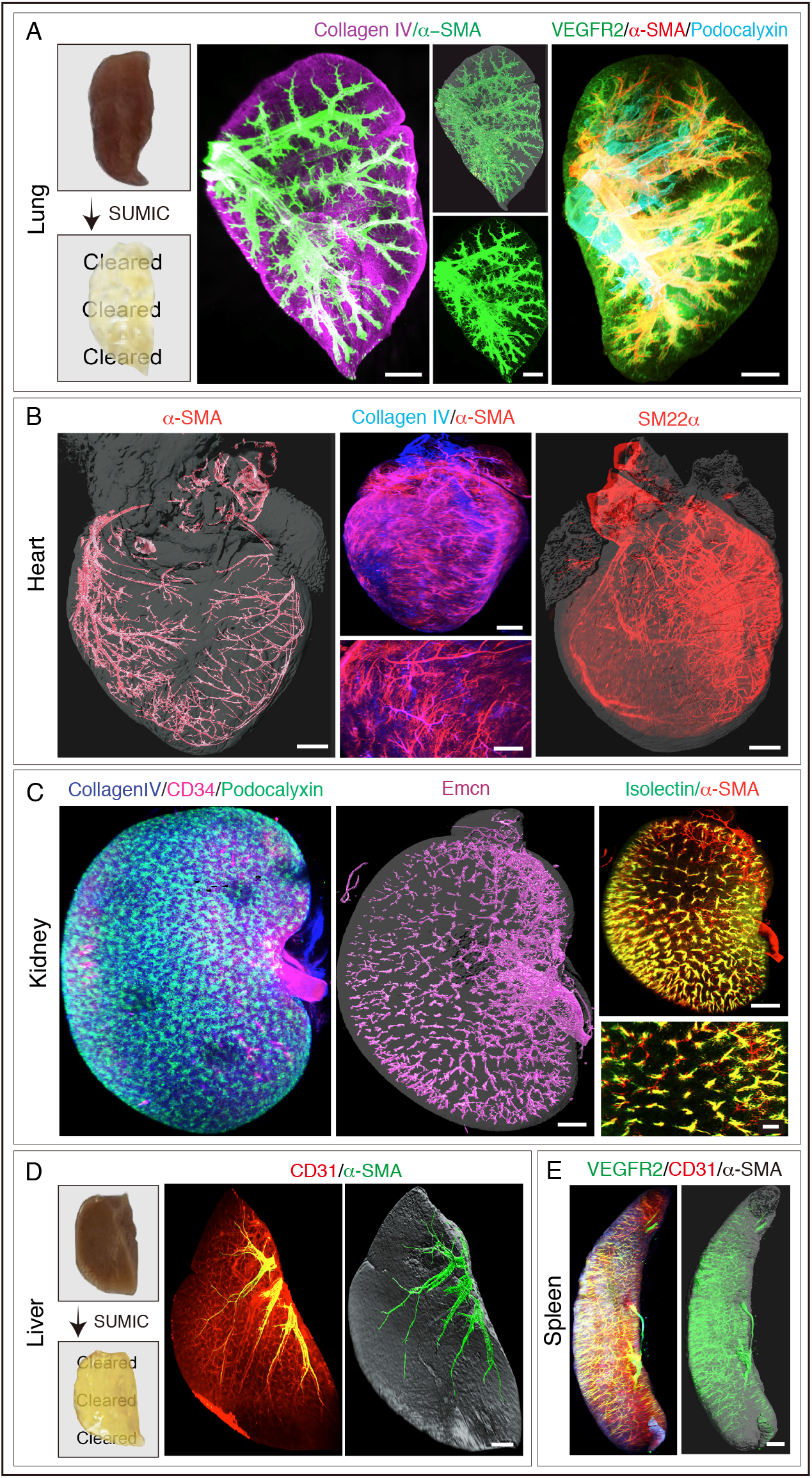
Light-sheet imaging of whole cleared organs using SUMIC. **A)** Photos (left) show mouse lung transparency before and after SUMIC. Representative images (middle) of whole cleared lung immuno-stained with Collagen IV and α-SMA acquired on a light sheet microscope. 3D image (right) of whole cleared lung immuno-stained with VEGFR2, α-SMA and Podocalyxin. **B)** Whole-organ imaging of cleared mouse heart stained with α-SMA (left), Collagen IV and α-SMA (middle) as well as SM22α (right). Inset shows high magnification of specific region. **C)** Light-sheet imaging of whole cleared mouse kidney staining as indicated. Inset shows high magnification of specific region. **D)** Photos (left) show mouse liver transparency before and after SUMIC. Representative 3D imaging (right) of whole cleared liver immuno-stained with CD31 and α-SMA. **E)** Representative 3D imaging of whole cleared spleen immuno-stained with VEGFR2, CD31 and α-SMA. Scale bars are 500 μm (A); Scale bars are 700 μm for whole-organ images and 200 μm for the high magnification insets (B); Scale bars are 500 μm (C-E).

**Fig. 3.**
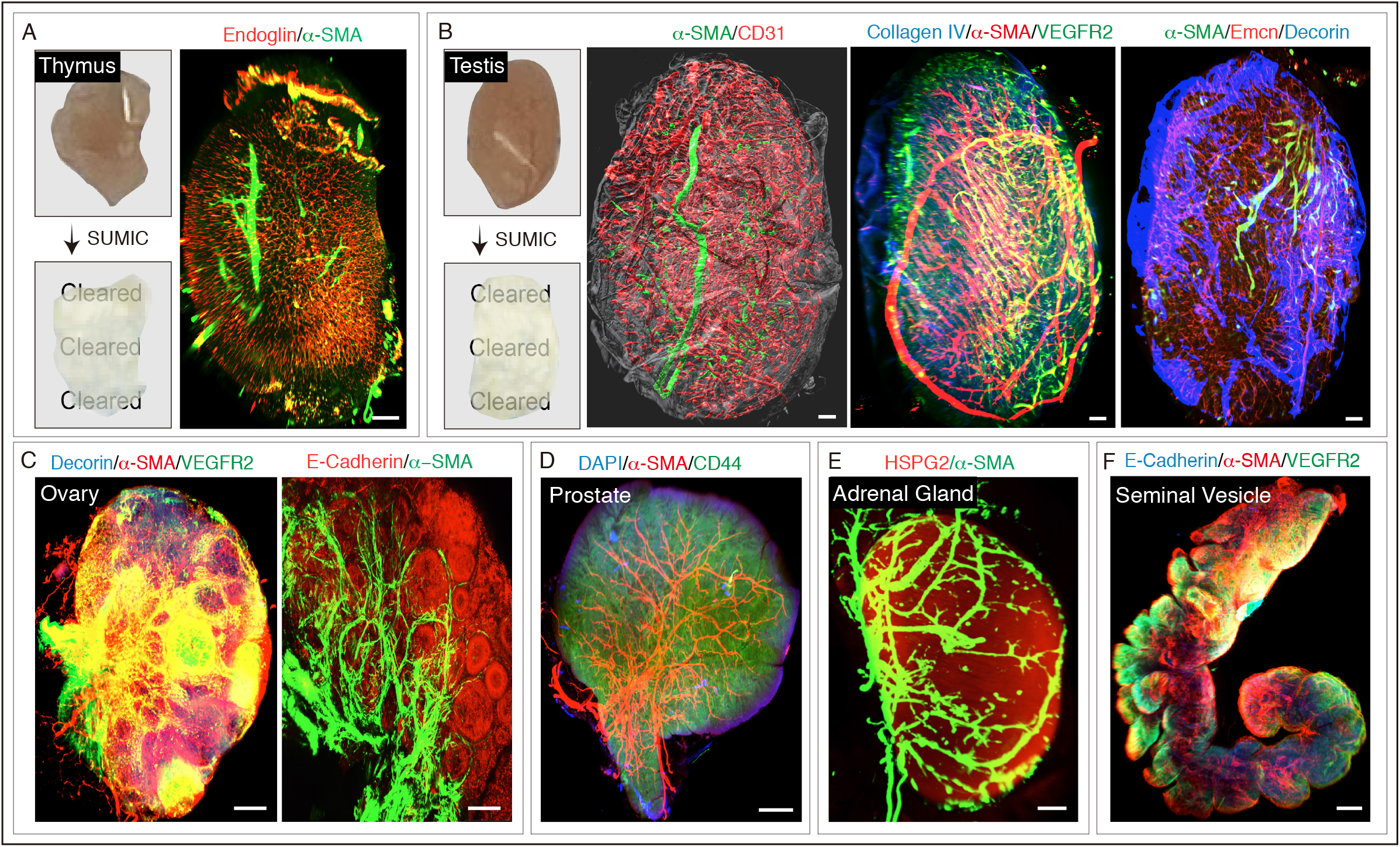
Light-sheet imaging of whole cleared endocrine glands using SUMIC. **A)** Photos (left) show mouse thymus transparency before and after SUMIC. Whole-organ 3D images (right) of cleared thymus stained with Endoglin and α-SMA acquired on a light sheet microscope. **B)** Photos (left) show mouse testis transparency before and after SUMIC. Representative 3D images (right) of whole cleared testis immunostained with the antibodies indicated in the panel. **C)** Whole-organ imaging of cleared mouse ovary stained with Decorin, α-SMA and VEGFR2 (left) as well as E-cadherin and α-SMA (right). **D)** 3D image of whole cleared prostate with α-SMA, CD44 and DAPI immunostaining.**E)** Light-sheet imaging of whole cleared adrenal gland immunostained with HSPG2 and α-SMA. **F)** Representative 3D imaging of whole cleared seminal vesicle stained with E-cadherin, α-SMA and VEGFR2. Scale bars are 400 μm (A-C and E); Scale bars are 500 μm (D); Scale bars are 900 μm (F).

**Fig. 4.**
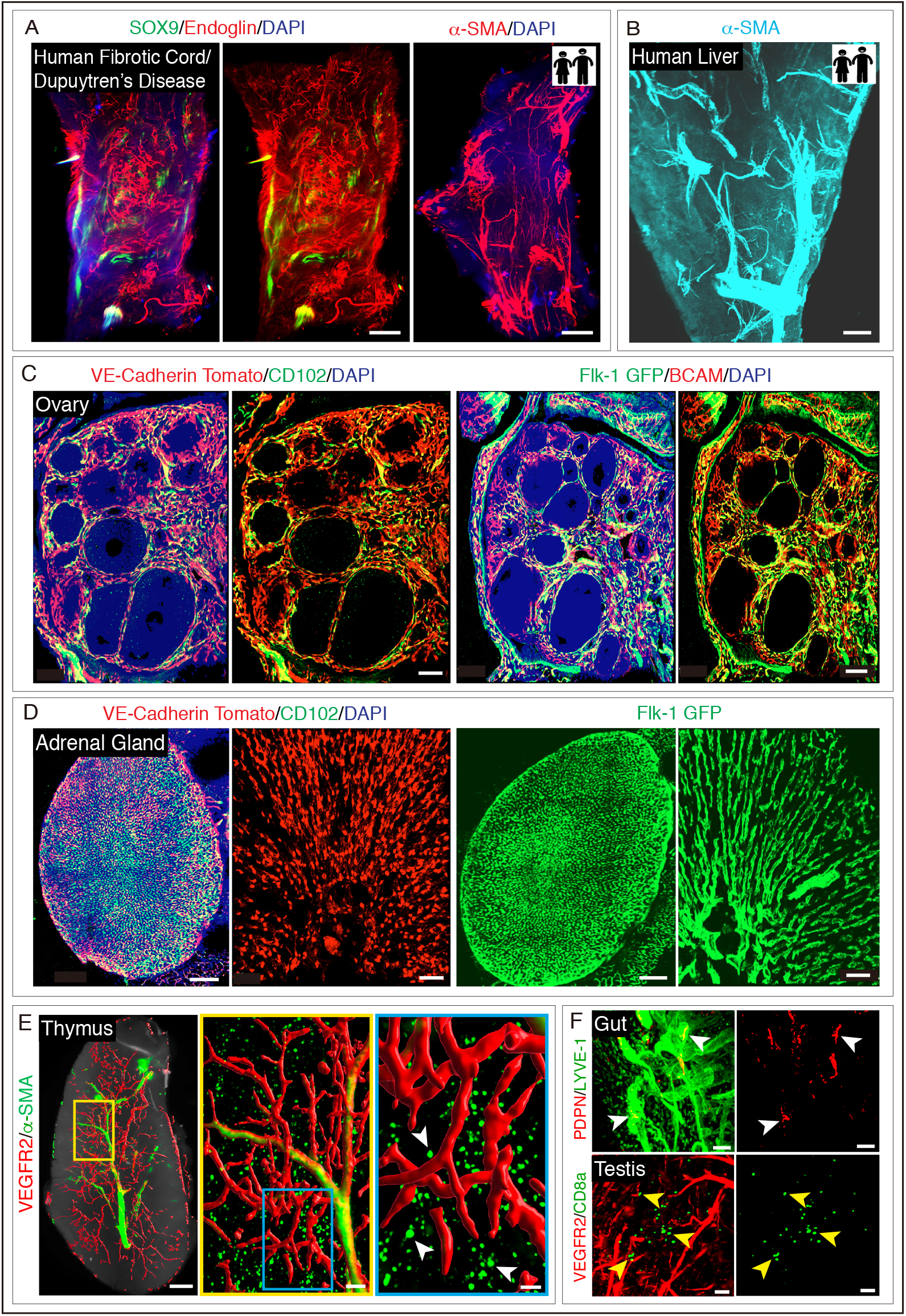
3D imaging of whole cleared endocrine glands and human tissues and detection of rare cells in whole organs using SUMIC. **A)** Representative 3D image of a cord of Dupuytren’s disease immuno-stained with SOX9 and Endoglin (left) as well as α-SMA (right). Nuclei: DAPI. **B)** Representative 3D imaging of the cleared human liver with α-SMA immunostaining. **C)** Representative 3D images (left) of cleared ovary stained with CD102 from *Cdh5(PAC)-CreERT2; Rosa26-td-Tomato* mutant mice acquired on a confocal microscope. 3D images (right) of cleared ovary immuno-stained with BCAM from Flk-1 GFP reporter mice. Nuclei: DAPI. **D)** Representative 3D images (left) of cleared adrenal gland stained with CD102 from *Cdh5(PAC)-CreERT2; Rosa26-td-Tomato* mice. 3D images (right) of cleared adrenal gland from Flk-1 GFP reporter mice. Nuclei: DAPI. **E)** Whole-organ imaging of cleared mouse thymus stained with VEGFR2 and α-SMA acquired on a light sheet microscope. Inset shows high magnification of specific region. Arrowheads indicate α-SMA^+^ stromal cells **F)** Representative 3D images (top) of cleared gut stained with Podoplanin (PDPN) and LYVE-1. Arrowheads (white) indicate Podoplanin^+^ lymphatic endothelial cells. 3D images (bottom) of cleared testis stained with VEGFR2 and CD8a. Arrowheads (yellow) indicate CD8a^+^ cells. Scale bar is 400 μm (A-B); Scale bars are 100 μm (C); Scale bars are 150 μm for tile scan images and 50 μm for the high magnification insets (D). Scale bars are 400 μm for whole-organ image, 100 μm and 50 μm for insets (E); Scale bars are 50 μm (F).

### SUMIC preserves tissue architecture and endogenous GFP/tomato fluorescence

To determine the effect of SUMIC processing on tissue morphology, we embedded SUMIC-processed organs in paraffin and examined the samples previously imaged in 3D by conventional histology. 2D histology showed that SUMIC processed tissues remained accessible to a conventional histological analysis by haematoxylin and eosin (Fig. S2). Tissue architecture remained unaffected, and different structures, such as blood vessels, were detected in SUMIC processed tissues similar to the control tissues processed by conventional paraffin embedding (Fig. S2). Thus, SUMIC enabled rapid, deep tissue immunolabeling and clearing of intact organs while maintaining tissue architecture.

Endogenous fluorescence loss is a crucial concern for solvent-based tissue clearing medium and other steps during tissue processing. To assess the impact of SUMIC clearing on endogenous fluorescence either *Flk1 GFP* or *CDh5 Cre ERT2 X td-tomato* mice was used. Our analysis revealed that the SUMIC processed samples preserved tomato and GFP fluorescence (Fig. 4, C and D).

### SUMIC permits single-cell resolution analysis in whole-organs

Light sheet microscopy was used to perform volumetric imaging and thereby produce the whole organ and tissue overviews of SUMIC-processed murine whole organs and human tissues. We next aimed at transitioning from a general overview of the complete tissue to the analysis of specific regions within the tissue at the single-cell resolution. Subsequently, computational surface rendering was applied to the regions of interest, which provides single-cell level information from three-dimensional tile scan images (Fig. 4, E and F). This analysis led to the detection of a small subset of round α-smooth muscle actin positive cells within the cortex region of the thymus in addition to the usual periarteriolar location of these cells (Fig. 4E). Such round α-SMA positive monocyte-macrophage subset has been reported in the bone marrow where these cells preserve primitive hematopoiesis (*36*). However, such a small subset of α-SMA positive monocytes-macrophages has not been reported to be present in the thymus.

### SUMIC enables quantitative imaging of whole organ overviews

Next, we carried out a quantitative analysis of the lymphatic vessels in 3D by LYVE-1 and Prox1 immunostainings on whole organs and endocrine glands. The circulatory system encompasses both the blood vascular and lymphatic vascular systems. The lymphatic system regulates fluid homeostasis, waste clearance and immune responses (*37, 38*). Recent studies have not only revealed age-dependent changes in blood vessels, organs and endocrine glands but also documented the correlation of these changes with the organ function (*39*). Here, we examined and compared the lymphatic vessels across different endocrine glands. Quantitative analysis of LYVE-1 and Prox1 immunostainings across endocrine glands from young and aged mice demonstrated expansion of lymphatic vessels in ovaries, testis and thyroid glands (Fig. 5, A and B). In contrast, the lymphatic vessel density remained unchanged across murine organs with age (Fig. S3).

**Fig. 5.**
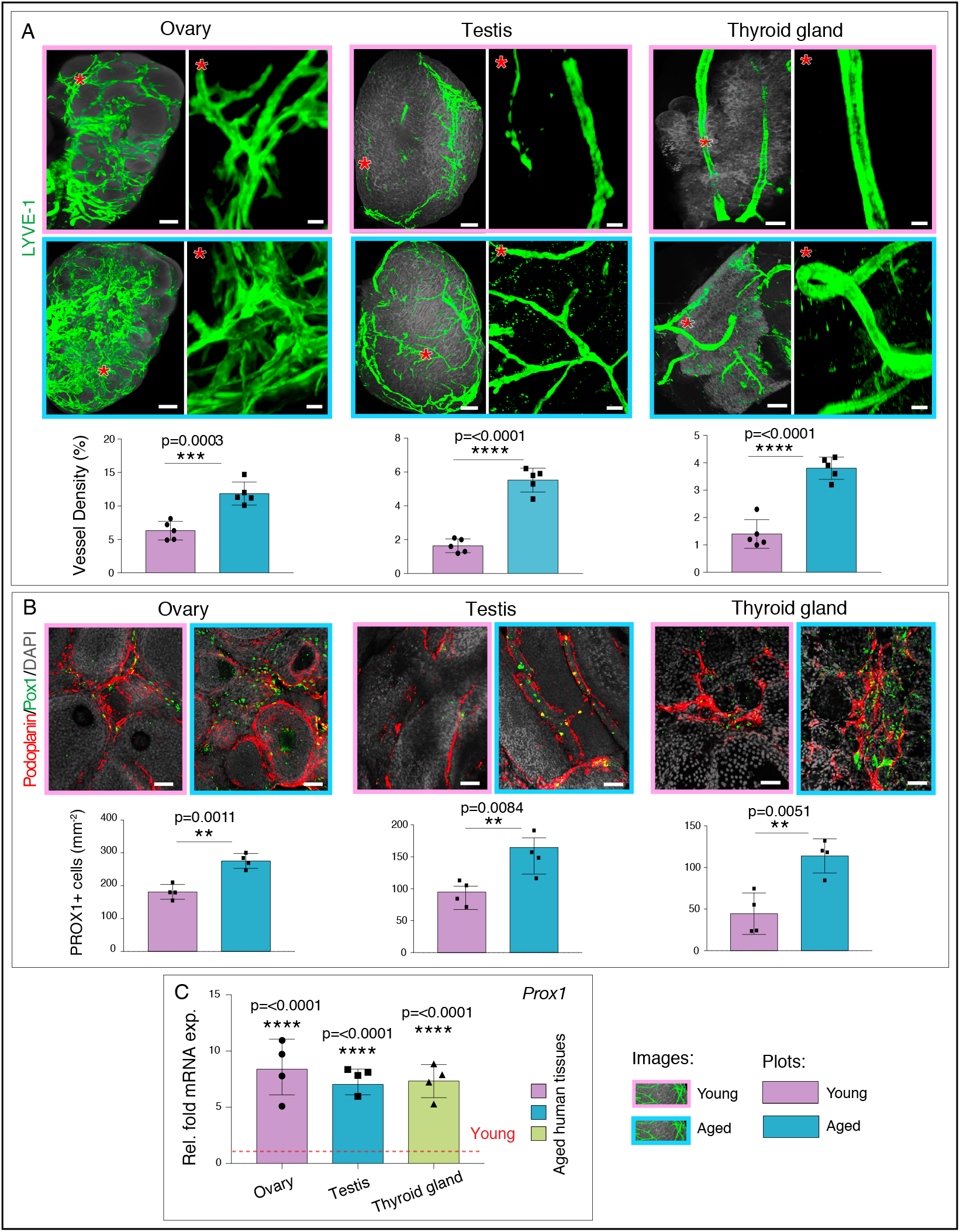
Age-associated lymphatic vessel declines in endocrine glands. **A)** Whole-organ imaging of cleared mouse ovary, testis and thyroid gland from young and aged mouse immuno-stained with LYVE-1. Asterisks indicate high magnification of specific areas in the different glands. Bar graphs show quantifications of lymphatic vessel density in young and aged ovary, testis and thyroid gland (*n=5*). **B)** 3D confocal images of young and aged ovary, testis and thyroid gland with Podoplanin and PROX1 immunostaining. Nuclei: DAPI. Bar graphs show quantifications of PROX1^+^ cell number in young and aged ovary, testis and thyroid gland (*n=4*). **C)** qPCR analysis of *Prox1* expression (normalized to *Actb*) in the aged human ovary, testis and thyroid compared to young endocrine glands (*n=5*). Data represents mean±s.d., p-value derived from two-tailed unpaired *t*-tests is given for all graphs. **: P <0.01; ***: P <0.001; ****; P <0.0001. Scale bars are 400 μm for whole-organ images and 50 μm for the high magnification insets (A); Scale bars are 50 μm (B).

To understand the relevance of age-related changes observed in these three murine endocrine glands, we analyzed human endocrine tissues. Healthy human testis, ovary and thyroid gland of young (<20 years) and aged (>70 years) individuals (Table S3) were tested for the *Prox1* gene by quantitative PCR (qPCR) (Fig. 5C). Likewise, murine endocrine glands, aged human ovary, testis and thyroid glands exhibited an increased *Prox1* expression (Fig. 5C). Importantly, analysis of even 4 random human samples of each of these glands showed these significant changes. These results provide evidence for the applicability of SUMIC method, which is not only rapid and robust but also holds tremendous potential in providing new insights and uncovering biological complexities.

## Discussion

3D organization of functional networks, such as blood or lymphatic vessels within organs and spatial information of cells within tissues, is important to understand the biological complexity (*10, 40–43*). 3D imaging of cleared large tissues or whole organs provides this information (*12, 14, 41, 43–45*). Further, it is a valuable alternative to traditional histology or immunofluorescence of thin tissue sections, both of which lack the crucial 3D information (*16, 45–48*). To enable rapid, simplified and high-resolution 3D imaging of whole organs and large tissue samples, we introduce and describe the SUMIC method. SUMIC enables the 3D molecular analysis of intact organs and tissue biopsies at a single-cell and subcellular resolution. A sequence of tissue alterations renders the samples susceptible to rapid and optimum immunostaining and histological staining while preserving cell and tissue integrity. Digestion of extracellular matrix by Collagenase and use of methanol increases antibody penetration. Collagenase based extracellular matrix digestion and incubation of antibodies at a higher temperature facilitate speed through rapid penetration of the antibodies. Further, these steps avoid the time-consuming delipidation steps that involve the use of highly toxic substances (*44, 49*). Further, the bleaching step reduces light scattering (*49, 50*). Hydrogen peroxide used for bleaching reduces tissue autofluorescence and is useful, particularly in tissues with high heme content, such as spleen and heart (*51*). Bleaching before the immunostaining aids in the retention of these fluorochromes which otherwise are susceptible to bleaching. Thus, SUMIC neither requires an injection of antibodies, nor it relies on transgenic mice expressing fluorescent reporters; it instead uses multicolor immunolabelling by fluorochrome-coupled antibodies.

ECi and PEGM have been used independently in various clearing methods as tissue clearing recipe components or for RI matching due to their inexpensive and flexible nature (*11, 51–53*). Here, we combined these two; ECi and PEGM, which facilitates the rapid clearing of larger tissues and organs. After clearing, images can be acquired on different imaging systems such as a laser scanning confocal microscope for single-cell and subcellular resolution or light sheet microscope for whole organ overviews. Therefore, SUMIC enables multicolor antibody-based immunolabelling, provides good tissue transparency, and preserves cellular morphology and flexibility in microscope usage.

As a result of facilitating high speed, high resolution and flexibility in organ, tissue, fluorochrome and microscope usage, SUMIC has the tremendous potential to make tissue clearing and 3D imaging more widely accessible to a broader audience. SUMIC is a versatile technique that can be used for answering a wide variety of questions throughout the biological sciences. Thus, SUMIC adds to the tools and information needed to investigate complex biological processes that demand spatial and architectural knowledge of cells within tissues and organs.

## Material and methods

### Mice

Organs and endocrine glands were dissected from adult 8-15-weeks-old male and female wild-type C57BL/6J mice purchased from Jackson Laboratory and bred in-house. To examine lymphatic vessels during ageing, young mice of 8-10 weeks-old and aged mice at 56-70-week-old age were used. For the study of native fluorescence, the *Kdr^tm2.1Jrt^/J* (*54*) (Stock No: 017006, Jackson Laboratory) and *ROSA26 td-Tomato* (Stock No: 007914, Jackson Laboratory) (*55*), were used. For genetic labelling of the vasculature, *Cdh5(PAC)-CreERT2* transgenic mice (*56*) were mated with *ROSA26 td-Tomato* reporters. All the animals utilized for the studies were housed in the University of Oxford. The experimental procedures on mice were performed following the UK Home Office Animal Care guidelines on the Operation of the Animals (Scientific Procedures) Act 1986 and the protocols approved by the local Animal Welfare and Ethical Review Board and by the UK Government Home Office (Animals Scientific Procedures Group).

### Tamoxifen treatment

For oral gavage of tamoxifen, tamoxifen (Sigma, T5648) was firstly dissolved in 100% ethanol and then suspended in corn oil to a final concentration of 5 mg/ml. For *Cdh5(PAC)-CreERT2; Rosa26-td-Tomato* transgenic mice, to induce Cre activity, mice were orally treated with tamoxifen at a dose of 50 mg/kg body weight for three consecutive days. Mice were sacrificed and analyzed two weeks after the last dose of tamoxifen.

### Tissue collection and preparation

Freshly dissected mouse tissues were rinsed in PBS (VWR, cat. No. 437117K) and then immediately transferred into ice-cold fixative solution of 4% paraformaldehyde (Sigma-Aldrich, P6148) and 0.05% glutaraldehyde (Sigma-Aldrich 340855) solution for 3 hours. The fixative should be freshly prepared and kept ice cold during its use. During the dissection of organs, the adjacent muscles and fat should be removed completely as their presence may subvert the successful generation of whole organ overviews during imaging. The samples were then washed with PBS for 3 times at room temperature on a shaker. Each wash lasted for 5 minutes. From our experience, the fixed samples can be stored for up to 4 days at 4°C before moving forward with the bleaching or antigen retrieval and permeabilization. While placing tissues in fixative works well for several murine organs, the perfusion fixation with 4% PFA is recommended to facilitate instant fixation to preserve the life-like state of the tissues.

The fixative is hazardous; highly flammable and toxic and must be handled with appropriate PPE and other safety measures.

### Bleaching

In larger heme rich organs bleaching was used to achieve maximum downstream organ clarity. Organs were immersed in a methanol gradient of 50%, 80% and 100% for 40 minutes each for dehydration. 100% methanol was changed twice, 20 minutes apart. These dehydration buffers should be ice-cold. Samples were then immersed in 5% (v/v) H_2_O_2_ in methanol for up to 3 hours (Sigma-Aldrich Cat No: H1009) for bleaching. Samples were then rehydrated with the reverse methanol concentrations. Following that, samples were washed three times with PBS for 20 minutes each. Dehydration for bleaching could also be achieved with Isopropanol that has a less toxic effect. However, methanol is better for removing pigmentation and more effective for delipidation. It also helps with membranes permeabilization and therefore antibody penetration.

### Antigen Retrieval, Permeabilization and Collagenase Digestion

The fixed samples were subjected to antigen retrieval, and permeabilization with the ice-cold buffer containing 25% urea (VWR, Cat. No. 28876.367), 15% glycerol (VWR, cat. no. 24388.260) and 7% Triton X-100 (Sigma-Aldrich, cat. no. T8787) diluted with ddH20 and incubated for 6 to12 hours at 4°C. Samples can then be directly processed to the next step of Collagenase based matrix digestion. Matrix digestion was performed with freshly prepared 0.2% Collagenase A (Merk, cat. no. 10103578001) prepared in PBS. Samples were incubated with Collagenase A solution for 30 minutes at 37°C on constant shaking. The Collagenase A solution should be freshly prepared. The samples were then washed twice for 5 minutes with the wash buffer containing 2% FBS (Sigma-Aldrich, cat. no. F7524) in PBS.

### Immunolabelling

For immunostaining, the organs were transferred to 5ml tubes and blocked in 10% donkey serum (Abcam, cat. no. ab7475), 10% DMSO with 0.5% Triton X-100 in PBS, at 37°C for 20 minutes. This blocking solution should be freshly prepared. Following blocking, the samples were incubated with the Alexa fluor conjugated antibodies or with the unconjugated primary antibodies. Primary antibody solution was prepared in 2% v/v donkey serum, 10% DMSO and 0.5% Triton-X-100 in PBS. The samples were incubated either overnight or for 14 to 16 hours at 37°C in a shaking water bath (Stuart, SBS40) rotating at 70 rpm with the primary antibodies (antibodies and dilution are listed in Table 1). Following incubation, the samples were washed with 2% v/v donkey serum, and 0.5% Triton X 100 in PBS for 3 hours at 37°C in the shaking water bath rotating at 70 rpm. The wash buffer was changed every 15 minutes for the first hour and then every 30 minutes for the remaining 2 hours. For the unconjugated primary antibodies; after the washing steps, the samples were incubated with the Alexa fluor coupled secondary antibodies for 6 to 8 hours at 37°C in the shaking water bath rotating at 70 rpm. Secondary antibody solution was prepared in 2% v/v donkey serum, 10% DMSO and 0.5% Triton-X-100 in PBS. Following incubation, the samples were washed with 2% v/v donkey serum, and 0.5% Triton X 100 in PBS for 3 hours at 37°C in the shaking water bath rotating at 70 rpm.

### Dehydration and tissue clearing

The immunostained samples were dehydrated in a gradient of isopropanol. Briefly, samples were immersed in a gradient of 30%, 50% and 80% (v/v) isopropanol for 30 minutes each under gentle rotation at room temperature. The organs were then immersed in 100% isopropanol for 1 hour with the 2 changes of isopropanol during this duration. The isopropanol was then completely removed, and the samples were rinsed twice with ethyl cinnamate (ECi) (Sigma-Aldrich, cat. no. 112372) for 5 minutes each. Samples were then cleared with 80% ECi and 20% poly(ethylene glycol) methyl ether methacrylate (PEGM) (Sigma, cat: 447943) under gentle rotation at room temperature for a minimum of 30 minutes.

### Human samples

Formalin-fixed paraffin-embedded (FFPE) normal testis, ovary and thyroid gland blocks were ordered from AMS Biotechnology (Oxford, UK). Human specimens provided by AMS Biotechnology (Europe) Limited are legally procured under the laws and regulations of the country. All samples are collected from consented patients in the major research/clinical centres under the local EC/IRB-approved protocols. Samples from donors at ages 18-20 years and 70-80 years were selected to compare the young and aged groups, respectively. Histological study was performed to confirm that sample tissues were healthy and lack disease components. Further details on the healthy human tissue samples are provided (Table S3).

### Imaging set-up, Image acquisition and Analysis

The cleared samples were imaged on the Miltenyi-LaVision Biotech UltraMicroscope II with the LaVision BioTech ImSpector software (MACs, Miltec bio). The microscope was equipped with a 2X objective lens for zoom body with a manual zoom of 0.63X– 6.3X. The objective was fitted with a Dipping Cap [5.7 mm] including correction Optics for Olympus MVPLAPO 2X. The microscope was fitted with the 405-100, 488-85, 561-100, 639-70 and 785-75 laser lines and the images captured with the Neo sCMOS camera (Andor). For image acquisition, cleared samples were manually attached with a drop of Superglue (No Nonsense, UK) to the sample holder adapter. Once the sample was firmly affixed to the sample holder, they were gently immersed in ECi in a quartz glass cuvette and excited with light sheets (30-90mm, dependent on organ size) of different wavelengths (488, 561, 640 and 785nm. The individual TIFF raw data images were converted by Imaris File converter (version 9.6.1, Bitplane) then analyzed by Imaris software (version 9.6.0, Bitplane).

Confocal immunofluorescent images of whole organs or regions of interesr were captured on a Zeiss Laser Scanning Microscope (LSM) 880 equipped with 7 laser lines (405, 453, 488, 514, 561, 594, and 633 nm), axio examiner (upright) stand and Colibri7 epifluorescence light source with LED illumination, the 10X Plan Apo 0.45 WD=2.0 M27 dry lens was used to image large regions using the tile scan function. Tiles were chosen based on organ size, and images were taken with a 10% overlap in order to stitch them together using Zen Black (version 3.1, Zeiss) software. Organs were imaged through the z-plane and reconstructed in Imaris in order to visualize the maximum intensity projection. In order to visualize the boundary of the organ, the autofluorescence from the 405 channel was converted into greyscale, and the 30% opaque image was manually overlayed with the corresponding TIFF file generated from Imaris. Imaris, Adobe Photoshop and Adobe illustrator software were used to generate, analyze and compile images.

### 3D surface rendering

To visualize components of the organ, 3D surface rendering was applied using the surface rendering tool in Imaris. First, the whole organ or region of interest (ROI) was defined and selected. Then, the smoothness of the resulting area was set up using the Smooth option. The threshold for background subtraction was manually annotated based on the individual image previewed.

### Quantifications of imaging data

To quantify the vascular density of whole organs, quantifications were performed using the Imaris Surface Analysis XTensions tool. To obtain total tissue volume, the 3D crop tool was used and analyzed with the Volume statistics function of Imaris. A single channel lymphatic marker was reconstructed using the 3D rendering surface function, and the tissue volume of the vessels was measured by employing the Surface statistics function in Imaris. The density of the lymphatic vasculature was calculated, dividing the tissue volume of vessels in the numerator by the total tissue volume in the denominator.

Quantification of PROX1^+^ cell number was done with Imaris software (version 9.2.1) based on single-cell resolution 3D images. Nuclei detection and membrane detection function in the Cell module of Imaris were used to automatically segment and analyze the numbers of PROX1^+^ cells. Under the Cell Creation wizard of Imaris, PROX1 channel was selected as a source channel for membrane-based detection, and DAPI channel was selected for nuclei-based detection. Nuclei locations were used as seed points for an algorithm performing a cell membrane calculation, which was used to distinguish between the inner and outer boundaries of the cell. The number of PROX1^+^ cells per tissue volume (mm^−3^) was calculated by dividing the number of segmented cells in the numerator by the total tissue volume in the denominator.

### Histology

Mice were sacrificed, and organs (liver, kidney, spleen and heart) were collected. One group of organs were fixed with 10% formalin in PBS. Another group of organs were fixed with 10% formation followed by the steps of SUMIC protocol (dehydration with gradient isopropanol and clearing with ECi). All samples were embedded in a paraffin block and sliced into 5 μm thick sections. Following H&E staining, the samples were imaged by bright-field microscopy (Olympus Co., Japan).

### Quantitative PCR

For the analysis of *Prox1* expression levels from FPPE blocks of human endocrine tissues, 25 μm sections were generated from FPPE blocks. After deparaffinization, FFPE sections were immersed into xylene and then into 100% ethanol twice, followed by air drying for 15 min. Quantitative PCR (qPCR) was performed using TaqMan gene expression assays on ABI PRISM 7900HT Sequence Detection System. RNA extraction kit (QIAGEN, 73504) was used to perform RNA extraction based on the manufacturer’s instructions. RNA samples were immediately processed for cDNA preparation using SuperScript IV First-Strand Synthesis System (Invitrogen, 18091200). The FAM-conjugated TaqMan probes were used along with TaqMan Gene Expression Master Mix (Applied Biosystems, 4369510). The relative expression level gene was normalized based on the *Actb*.

### Statistical analysis

All statistical analyses were performed using GraphPad Prism software (version 9.1). All data are presented as mean± s.d. The significance of the difference in mean values was determined using two-tailed Student’s t-test. The statistical significance was declared when P <0.05. ns: not significant; *: P <0.05; **: P <0.01; ****: P <0.001; ****: P <0.0001. No randomization or blinding was used, and no animals were excluded from analyses. Several independent experiments were performed to guarantee reproducibility of findings.

## Supporting information

SUMIC processed lung showing multicolor immunostaining for endothelial and perivascular cell markers.

## Acknowledgements

We thank Dr. Lynn Williams and Dr. Nan Yang for their help in providing human tissue samples. A.P.K is supported by Medical Research Council (CDA: MR/P02209X/1), European Research Council (StG: metaNiche, 805201), Leuka (2017/JGF/001), The Royal Society (RG170326), Kennedy Trust for Rheumatology Research (KENN 15 16 09), CRUK Development Fund (CRUKDF 0317-AK) and John Fell OUP Research Fund (161/061). This work is also supported by the Sir Henry Dale Fellowship (202300/Z/16/Z) from the Wellcome Trust and the Royal Society to S.K.R; and the Wellcome Trust PRF (100262Z/12/Z) and a KTRR grant to M.L.D. J.C is supported by the National Natural Science Foundation of China (81901060) and Science & Technology Key Research and Development Program of Sichuan Province (2019YFS0142).

## Author Contributions

L.B., J.C., J.D.A., A.C., S.K.R, and A.P.K. designed and organized the experiments. L.B., and J.C. performed most of the experiments. L.B., J.C., and A.P.K. analyzed data and interpreted results. J.N. and S.K.R. provided critical samples. M.L.D. contributed to the design of imaging setup, provided critical advice and commented on the manuscript. J.C. prepared figures. J.C and A.P.K wrote the paper. L.B. wrote methods section. J.C., J.D.A., A.C. and A.P.K. edited the manuscript. A.P.K. conceived, devised and supervised the study and method development.

## Author Information

The authors do not declare competing financial interests.

## Legends to Supplementary Figures

**Fig. S1.**
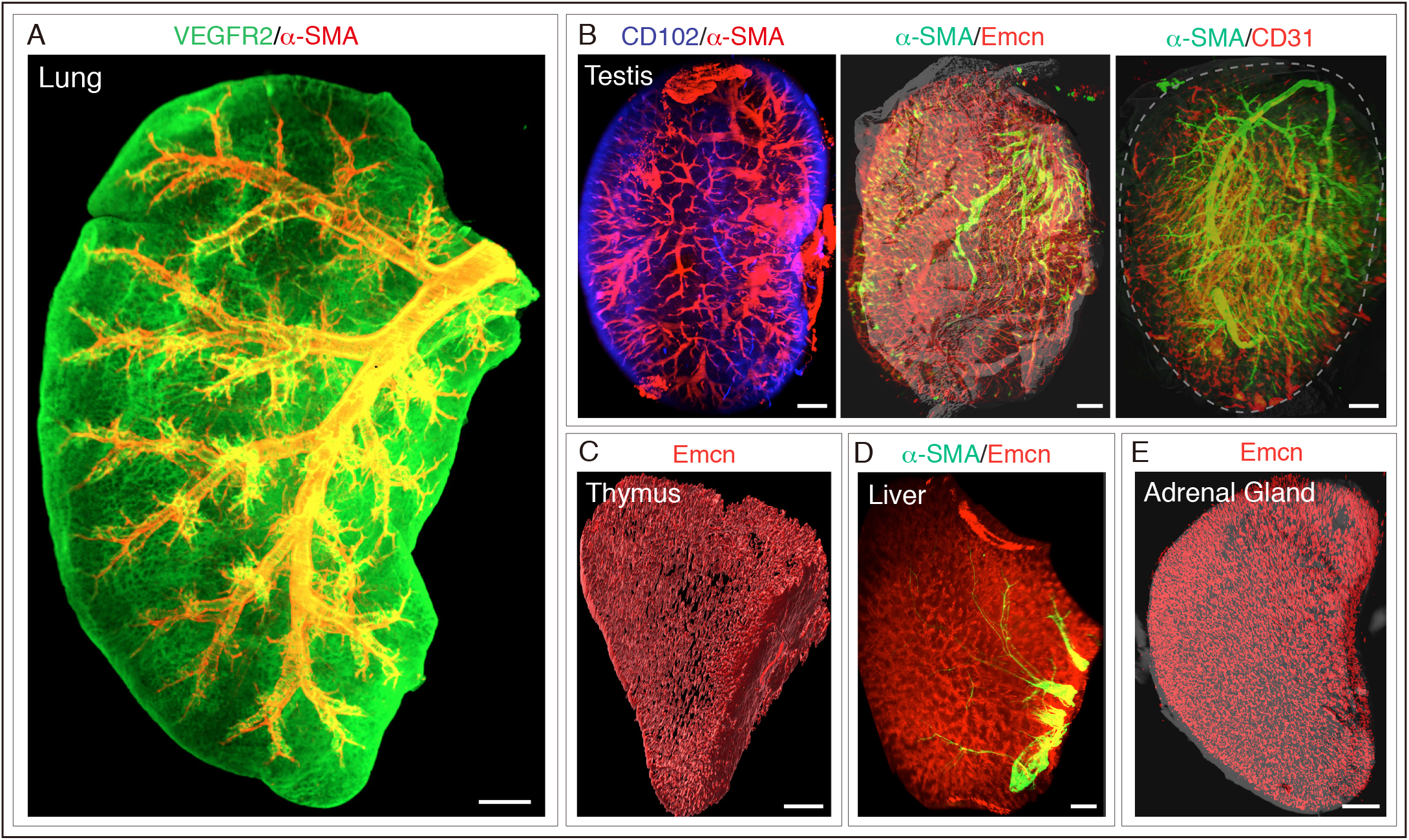
Light-sheet imaging of whole cleared mouse organs and endocrine glands. **A)** Light-sheet imaging of whole cleared mouse lung with VEGFR2 and α-SMA immunostaining. **B)** Representative 3D images acquired on a light sheet microscope of whole cleared testis stained with the antibodies indicated in the panel. **C)** 3D image of whole cleared thymus stained with Emcn. **D)** Whole-organ imaging of cleared mouse liver stained with α-SMA and Emcn. **E)** Whole-organ imaging of cleared adrenal gland stained with Emcn. Scale bars are 500 μm (A and D); Scale bars are 400 μm (B, C and E).

**Fig. S2.**
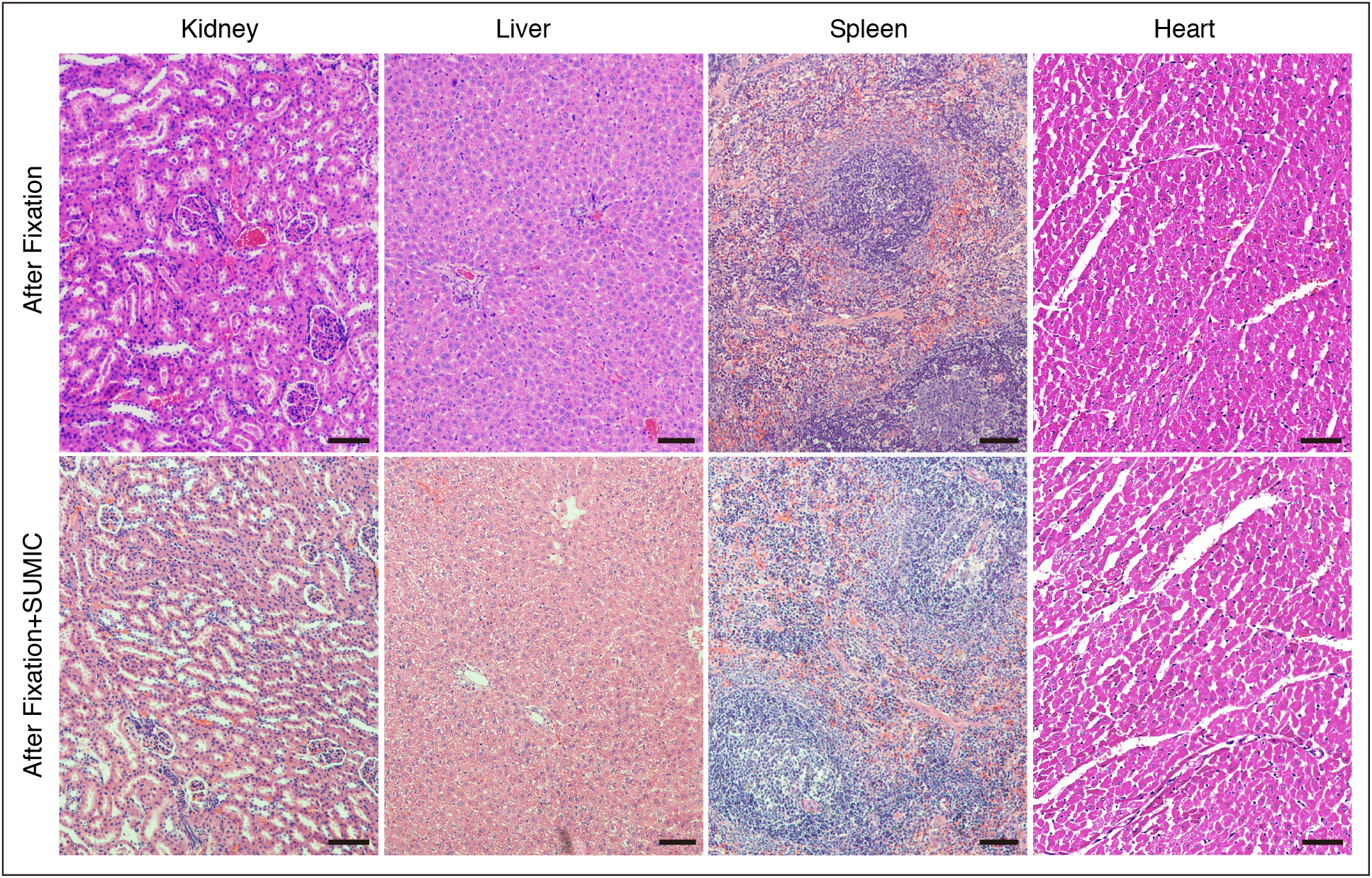
SUMIC preserves tissue integrity, and the tissues can be used for histology. H&E staining of mouse kidney, liver, spleen and heart after fixation (up) and fixation followed by SUMIC processes (below). Scale bars are 200 μm.

**Fig. S3.**
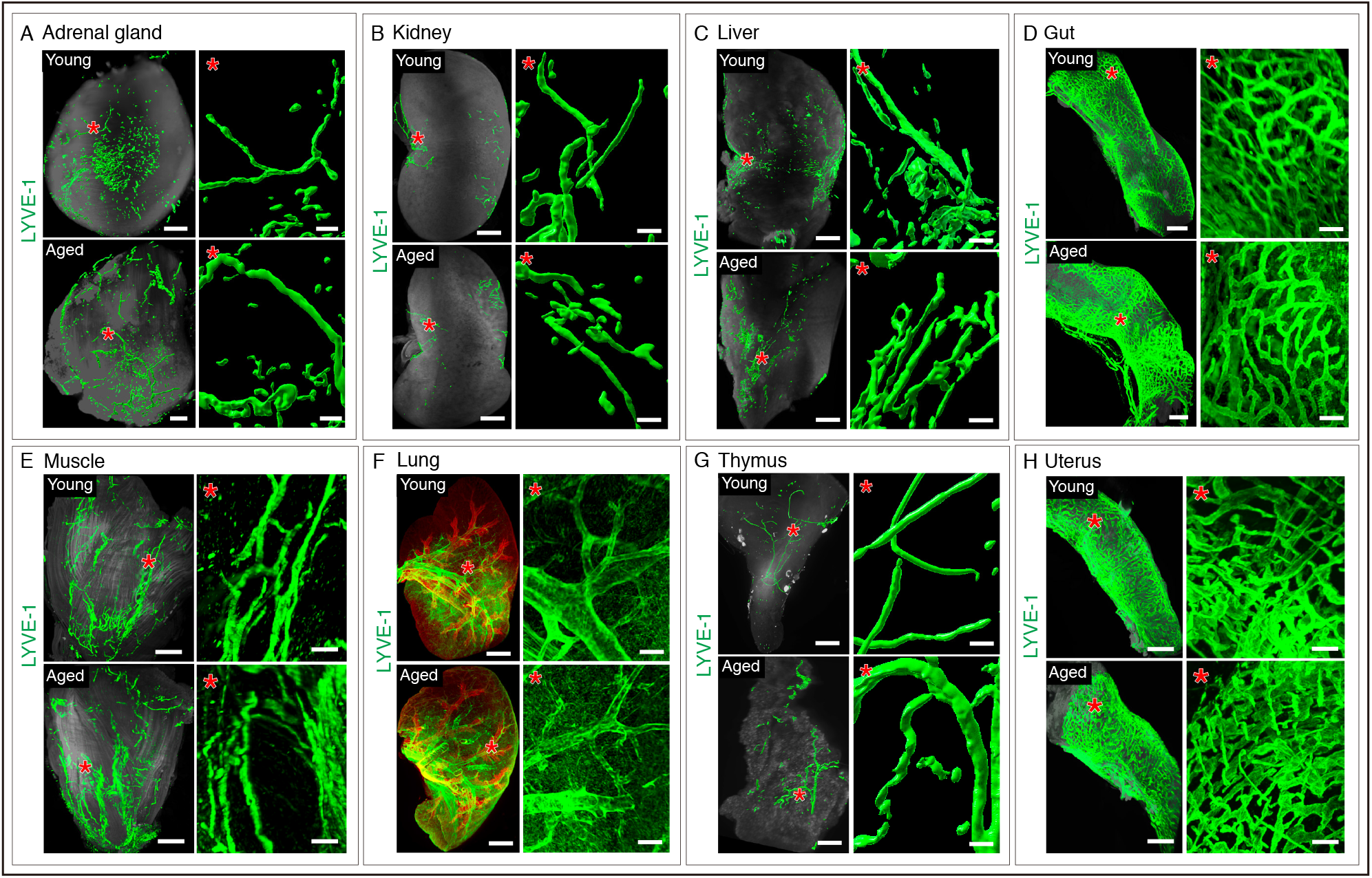
Lymphatic vessel imaging in young versus aged murine organs. **A-H)** Whole-organ imaging of cleared mouse adrenal gland, kidney, liver, gut, muscle, lung, thymus and uterus with LYVE-1 immunostaining. Asterisks indicate high magnification of specific areas in the different organs. Scale bars are 400 μm for whole-organ images and 50 μm for the high magnification insets (A-H).

## Legends to Supplementary Tables

**Table S1.**
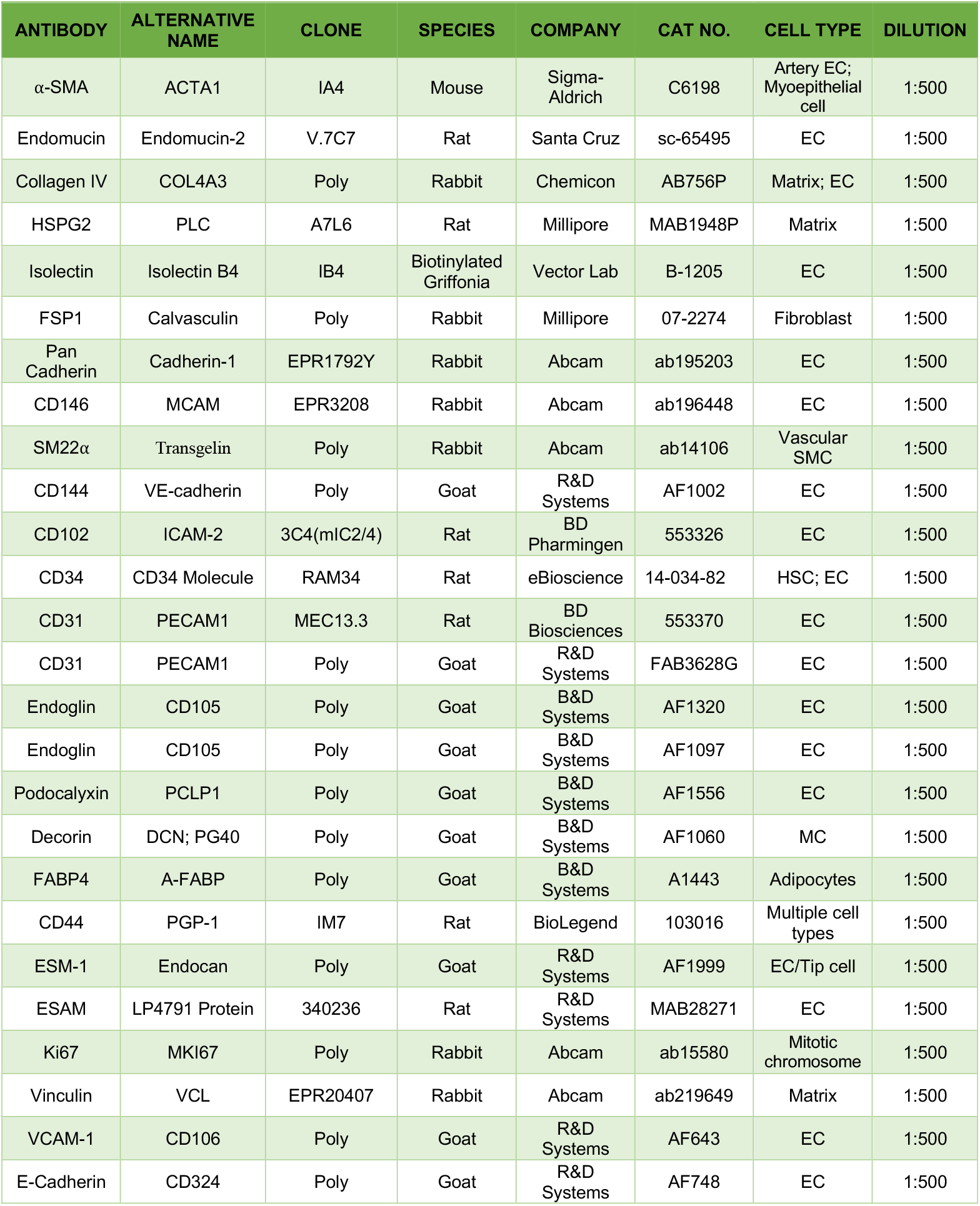
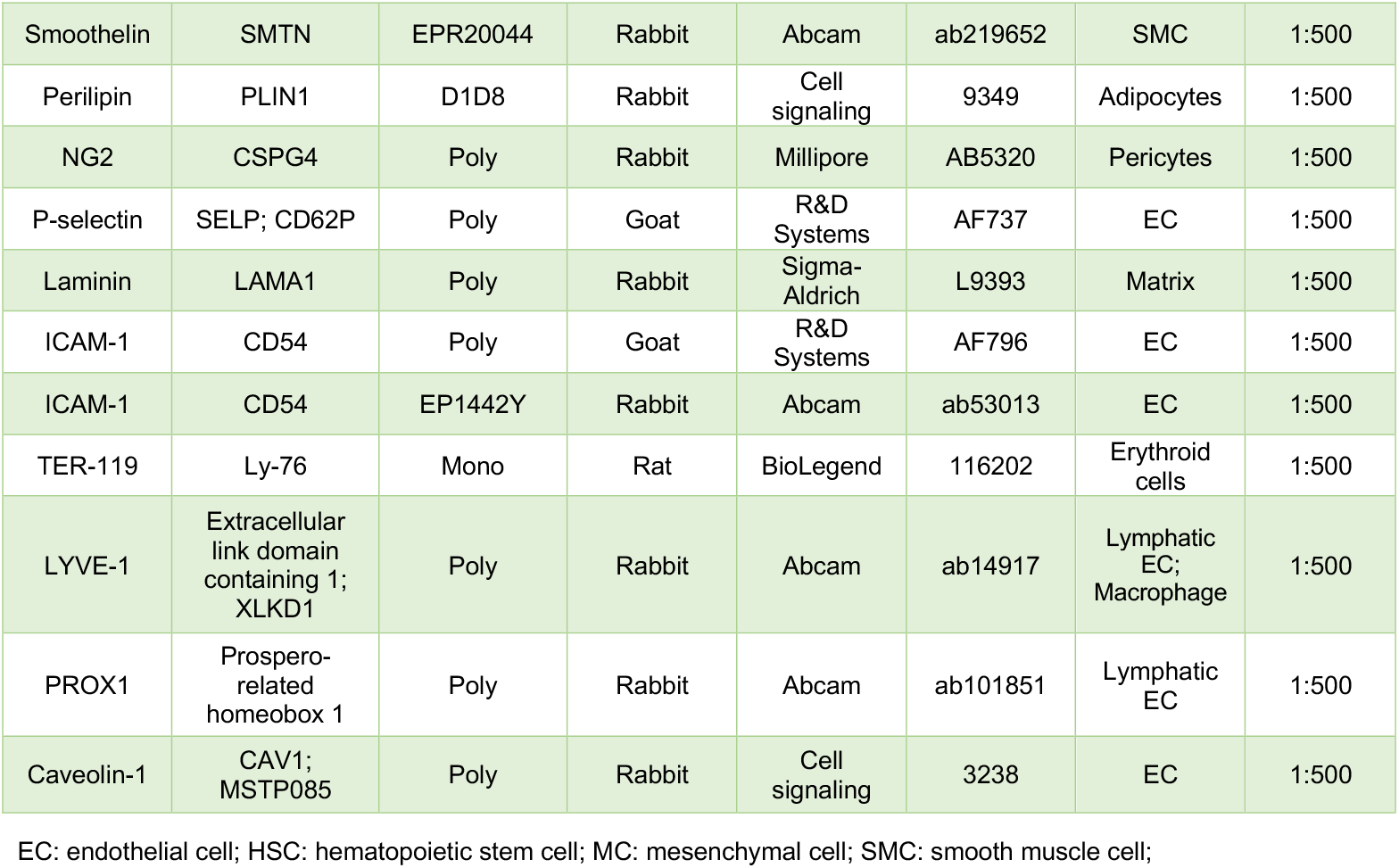
List of primary antibodies thoroughly tested with SUMIC

| ANTIBODY | ALTERNATIVE NAME | CLONE | SPECIES | COMPANY | CAT NO. | CELL TYPE | DILUTION |
| --- | --- | --- | --- | --- | --- | --- | --- |
| α-SMA | ACTA1 | IA4 | Mouse | Sigma-Aldrich | C6198 | Artery EC; Myoepithelial cell | 1:500 |
| Endomucin | Endomucin-2 | V.7C7 | Rat | Santa Cruz | sc-65495 | EC | 1:500 |
| Collagen IV | COL4A3 | Poly | Rabbit | Chemicon | AB756P | Matrix; EC | 1:500 |
| HSPG2 | PLC | A7L6 | Rat | Millipore | MAB1948P | Matrix | 1:500 |
| Isolectin | Isolectin B4 | IB4 | Biotinylated Griffonia | Vector Lab | B-1205 | EC | 1:500 |
| FSP1 | Calvasculin | Poly | Rabbit | Millipore | 07-2274 | Fibroblast | 1:500 |
| Pan Cadherin | Cadherin-1 | EPR1792Y | Rabbit | Abcam | ab195203 | EC | 1:500 |
| CD146 | MCAM | EPR3208 | Rabbit | Abcam | ab196448 | EC | 1:500 |
| SM22α | Transgelin | Poly | Rabbit | Abcam | ab14106 | Vascular SMC | 1:500 |
| CD144 | VE-cadherin | Poly | Goat | R&D Systems | AF1002 | EC | 1:500 |
| CD102 | ICAM-2 | 3C4(mIC2/4) | Rat | BD Pharmingen | 553326 | EC | 1:500 |
| CD34 | CD34 Molecule | RAM34 | Rat | eBioscience | 14-034-82 | HSC; EC | 1:500 |
| CD31 | PECAM1 | MEC13.3 | Rat | BD Biosciences | 553370 | EC | 1:500 |
| CD31 | PECAM1 | Poly | Goat | R&D Systems | FAB3628G | EC | 1:500 |
| Endoglin | CD105 | Poly | Goat | B&D Systems | AF1320 | EC | 1:500 |
| Endoglin | CD105 | Poly | Goat | B&D Systems | AF1097 | EC | 1:500 |
| Podocalyxin | PCLP1 | Poly | Goat | B&D Systems | AF1556 | EC | 1:500 |
| Decorin | DCN; PG40 | Poly | Goat | B&D Systems | AF1060 | MC | 1:500 |
| FABP4 | A-FABP | Poly | Goat | B&D Systems | A1443 | Adipocytes | 1:500 |
| CD44 | PGP-1 | IM7 | Rat | BioLegend | 103016 | Multiple cell types | 1:500 |
| ESM-1 | Endocan | Poly | Goat | R&D Systems | AF1999 | EC/Tip cell | 1:500 |
| ESAM | LP4791 Protein | 340236 | Rat | R&D Systems | MAB28271 | EC | 1:500 |
| Ki67 | MKI67 | Poly | Rabbit | Abcam | ab15580 | Mitotic chromosome | 1:500 |
| Vinculin | VCL | EPR20407 | Rabbit | Abcam | ab219649 | Matrix | 1:500 |
| VCAM-1 | CD106 | Poly | Goat | R&D Systems | AF643 | EC | 1:500 |
| E-Cadherin | CD324 | Poly | Goat | R&D Systems | AF748 | EC | 1:500 |
| Smoothelin | SMTN | EPR20044 | Rabbit | Abcam | ab219652 | SMC | 1:500 |
| Perilipin | PLIN1 | D1D8 | Rabbit | Cell signaling | 9349 | Adipocytes | 1:500 |
| NG2 | CSPG4 | Poly | Rabbit | Millipore | AB5320 | Pericytes | 1:500 |
| P-selectin | SELP; CD62P | Poly | Goat | R&D Systems | AF737 | EC | 1:500 |
| Laminin | LAMA1 | Poly | Rabbit | Sigma-Aldrich | L9393 | Matrix | 1:500 |
| ICAM-1 | CD54 | Poly | Goat | R&D Systems | AF796 | EC | 1:500 |
| ICAM-1 | CD54 | EP1442Y | Rabbit | Abcam | ab53013 | EC | 1:500 |
| TER-119 | Ly-76 | Mono | Rat | BioLegend | 116202 | Erythroid cells | 1:500 |
| LYVE-1 | Extracellular link domain containing 1; XLKD1 | Poly | Rabbit | Abcam | ab14917 | Lymphatic EC; Macrophage | 1:500 |
| PROX1 | Prospero-related homeobox 1 | Poly | Rabbit | Abcam | ab101851 | Lymphatic EC | 1:500 |
| Caveolin-1 | CAV1; MSTP085 | Poly | Rabbit | Cell signaling | 3238 | EC | 1:500 |

EC: endothelial cell; HSC: hematopoietic stem cell; MC: mesenchymal cell; SMC: smooth muscle cell;

**Table S2.** Troubleshooting table

| Problems | Possible reason | Solution |
| --- | --- | --- |
| Weak antibody signal | Antibody concentration is inadequate to stain the whole organ | Optimize the antibody concentration |
| Noisy and non-transparent fluorescence signal | The dehydration may be limited. It also affects tissue clearing | Ensure the dehydration is enough to get transparent signal as per protocol |
| Antibody precipitations on the organ | Inadequate washing | Ensure adequate wash is carried out to overcome this issue |
| Appearance of nonspecific fluorescence signal | Precipitates and aggregates in the secondary antibodies | Avoid freeze-thaw cycles and the antibody should be spun down and mixed gently before use |
| High background along with antibody signal | Too much pigmentation and Hematoma on samples; Inadequate washing steps | Bleaching required to remove pigmentation and reduce hematoma signal; Wash more times |
| Fade antibody staining | Dehydration of chemicals | Optimised dehydration as suggested on the protocol |
| Tissue clearing and preservation of fluorescence signal | Slow tissue clearing and loss of fluorescence signal within few days | PEG increases the hydration capacity of the clearing medium and preserve longer fluorescence signal |
| Sometimes bubbles build up inside the whole organ | Hole in the organ and last hydration step | Fill the empty space in the organ with clearing solution by needle without damaging the tissues |
| Unstained or dark regions toward the centre to the organ/tissue | Collagenase digestion was either insufficient or unsuccessful | Collagenase solution should be freshly prepared; collagenase should be incubated at 37°C with constant shaking |

**Table S3.** Details of healthy young and aged human endocrine gland tissue samples

| Serial Number | Sample Name | Sample AMSBIO ID | Catalogue number | Age | Gender | Diagnosis | Ethnicity | Year of Procurement |
| --- | --- | --- | --- | --- | --- | --- | --- | --- |
| 1 | Ovary | 4765 | AMS-33028 | 19 | female | normal | Caucasian | 09/11/2016 |
| 2 | Ovary | 5514 | AMS-33028 | 18 | female | normal | Caucasian | 19/07/2015 |
| 3 | Ovary | 6379 | AMS-33028 | 18 | female | normal | Caucasian | 03/12/2016 |
| 4 | Ovary | 7456 | AMS-33028 | 19 | female | normal | Caucasian | 15/05/2017 |
| 5 | Ovary | 2108 | AMS-33028 | 80 | female | normal | Caucasian | 30/07/2017 |
| 6 | Ovary | 1582 | AMS-33028 | 79 | female | normal | Caucasian | 13/03/2017 |
| 7 | Ovary | 6715 | AMS-33028 | 70 | female | normal | Caucasian | 12/01/2015 |
| 8 | Ovary | 4517 | AMS-33028 | 72 | female | normal | Caucasian | 28/08/2016 |
| 9 | Testis | 149 | AMS45022 | 18 | male | normal | Caucasian | 19/04/2016 |
| 10 | Testis | 152 | AMS45022 | 19 | male | normal | Caucasian | 03/05/2017 |
| 11 | Testis | 154 | AMS45022 | 18 | male | normal | Caucasian | 25/05/2017 |
| 12 | Testis | 156 | AMS45022 | 20 | male | normal | Caucasian | 11/09/2018 |
| 13 | Testis | 29300 | AMS45022 | 70 | male | normal | Caucasian | 22/09/2014 |
| 14 | Testis | 3509 | AMS45022 | 71 | male | normal | Caucasian | 01/07/2014 |
| 15 | Testis | 3077 | AMS45022 | 76 | male | normal | Caucasian | 27/08/2017 |
| 16 | Testis | 7267 | AMS45022 | 75 | male | normal | Caucasian | 18/03/2017 |
| 17 | Thyroid gland | 0235 | AMS-47028 | 18 | female | normal | Caucasian | 27/09/2015 |
| 18 | Thyroid gland | 3555 | AMS-47028 | 19 | female | normal | Caucasian | 19/06/2015 |
| 19 | Thyroid gland | 5279 | AMS-47028 | 19 | female | normal | Caucasian | 20/04/2015 |
| 20 | Thyroid gland | 6022 | AMS-47028 | 20 | female | normal | Caucasian | 03/10/2016 |
| 21 | Thyroid gland | 4274 | AMS-47028 | 71 | female | normal | Caucasian | 20/05/2016 |
| 22 | Thyroid gland | 3712 | AMS-47028 | 73 | male | normal | Caucasian | 01/12/2015 |
| 23 | Thyroid gland | 2640 | AMS-47028 | 70 | female | normal | Caucasian | 23/01/2016 |
| 24 | Thyroid gland | 4888 | AMS-47028 | 72 | female | normal | Caucasian | 17/02/2017 |

## Legends to Movies

**Movie 1.** SUMIC processed lung showing multicolor immunostaining for endothelial and perivascular cell markers.

**Movie 2.** SUMIC processed testis showing multicolor immunostaining for endothelial and perivascular cell markers.

**Movie 3.** SUMIC processed spleen showing multicolor immunostaining for endothelial and perivascular cell markers.

**Movie 4.** SUMIC processed thymus showing multicolor immunostaining for endothelial and perivascular cell markers.

**Movie 5.** SUMIC processed heart showing immunostaining for arteries.

**Movie 6.** SUMIC processed uterus showing immunostaining for a lymphatic marker.

